# From Attention Control to Stimulus Selection: Neural Mechanisms Revealed by Multivariate Pattern and Functional Connectivity Analyses

**DOI:** 10.64898/2026.04.10.717841

**Authors:** Qiang Yang, Sreenivasan Meyyappan, George R Mangun, Mingzhou Ding

## Abstract

Visual spatial attention deployed in advance of sensory stimulation enhances the processing of the stimuli at an attended location. While it is understood that the attention control signals are established even before the stimulus occurs, how these signals help achieve stimulus selection is still not clear. Here, we investigated the neural mechanisms of spatial attention control and subsequent stimulus selection by recording fMRI data from participants performing a cued visual spatial attention task. At the beginning of each trial, participants were cued to covertly attend either the left or the right visual field. Following a random cue-target period, a target stimulus appeared either at the attended location or at the unattended location. Participants discriminated the stimulus appearing at the attended location and ignored the stimulus appearing at the unattended location. Using MVPA decoding and multivariate functional connectivity techniques, we investigated the nature of the information in visual cortex during both the cue and target periods, and further probed how cue-related information was related to target-related sensory processing. The following results were found: (1) attend-left vs. attend-right conditions could be decoded from the cue-period neural activity in visual cortex, (2) target position (target-left vs. target-right) could also be decoded from the target-evoked activity in visual cortex, (3) classifiers built on cue-period neural activity could cross-decode attended target-evoked neural activity in visual cortex and vice versa, (4) higher pattern similarity across cue and target periods, as indexed by cross-decoding accuracy, was correlated with better behavioral performance, (5) the strength of cue-evoked multivariate functional connectivity patterns in visual cortex was positively correlated with behavioral performance, and (6) cue-evoked multivariate functional connectivity patterns were similar to those evoked by the attended targets, and higher connectivity pattern similarity across cue and target periods was correlated with better behavioral performance. These results suggest that top-down attention control enables the formation of (1) a spatial attention template at the level of individual visual cortical areas and (2) an attention network template across visual areas, and these neural patterns support stimulus selection likely via a template matching mechanism at both area and pathway levels.

## Introduction

Visual attention can be deployed voluntarily in anticipation of sensory input, in line with behavioral goals. It is generally accepted that following an attention-directing cue, top-down signals are generated within the frontal-parietal dorsal attention network (DAN), which then propagate to the visual cortex to bias neural activity, enhancing task-relevant information and suppressing task-irrelevant distractors. Extensive research has been carried out in both humans and nonhuman primates to test this model (S. L. Bressler et al., 2008; Corbetta et al., 2000; Giesbrecht et al., 2006; Hopfinger et al., 2000; Kastner et al., 1999; Luck et al., 1997; McMains et al., 2007; Serences et al., 2004; Sylvester et al., 2007). In a recent study, applying MVPA methods to fMRI data recorded in a cued visual spatial attention paradigm, we showed that in the cue-target period (anticipatory attention), top-down biasing signals modulated neural activity patterns in multiple retinotopic visual areas from primary visual cortex (V1) through higher-order areas, and the strength of these signals predicted behavioral performance (Meyyappan et al., 2025). Here we ask how these anticipatory neural signals interact with the incoming target stimulus to enable the selection of task-relevant information and enhance behavioral performance.

In feature-based or object-based attention, numerous functional magnetic resonance imaging (fMRI) studies have provided evidence that top-down anticipatory attention generates a neural pattern in visual cortex that resembles that observed during the perception of the target stimulus (Gayet & Peelen, 2022; Peelen & Kastner, 2011; Stokes et al., 2009). This anticipatory neural pattern, referred to as the attention template, is thought to select the sensory input via a template matching mechanism (Peelen & Kastner, 2011; Stokes et al., 2009), where the stronger the similarity between the attention template and the sensory input, the more effective the stimulus selection, and the better the behavioral performance. To what extent the attention template idea applies to visual spatial attention remains to be better understood. Recently, using color and location cueing in a visual search task, Jimenez et al. (2024) applied behavioral and ERP techniques to suggest that a location template was activated by the location cue before the onset of the search array, and the location-based attention template guides behavior as efficiently as the feature-based (color) template (Jimenez et al., 2024). To understand the underlying neural mechanisms, the first goal of this study was to test the hypothesis that the sensory biasing patterns in visual cortex activated by anticipatory attention to a spatial location resemble those evoked by an attended target stimulus appearing at the location, and the subsequent stimulus selection is achieved through template matching.

Upon the presentation of a visual stimulus, volleys of neural activity initiated in the retina course up multiple parallel channels, starting with the lateral geniculate nucleus (LGN), through cortical areas V1 and V2, to reach additional tiers of visual cortex in the dorsal (parietal lobe) and ventral (temporal lobe) pathways where the meaning of the stimulus is extracted and the appropriate behavioral response initiated (Mishkin et al., 1983). Attention, in addition to enhancing neural responses in individual visual areas, is also expected to enhance neural transmissions across visual areas, prioritizing the processing pathways relevant for behavioral goals. To what extent and for what purpose top-down visual spatial attention facilitates neural transmission pathways in advance of stimulus processing remains to be better understood. We postulate that a cue-evoked functional connectivity pattern, henceforth referred to as an *attention network template*, resembles that evoked by the target stimulus and increased similarity between the target-evoked functional connectivity pattern and the attention network template predicts better behavior. That is, at the network level, stimulus selection is achieved through network template matching. To test this network model of attention control, the univariate functional connectivity (UFC) between regions of interest (ROIs) in visual cortex is likely to be inadequate because it uses neural responses averaged across a ROI, thus ignoring pattern level information within visual areas. A novel approach, referred to as *multivariate functional connectivity* (MFC) here, has become available, which measures coordinated neural activities at the neural pattern level between different visual areas (Coutanche & Thompson-Schill, 2013). The second goal of this study is to apply MFC to test the attention network template hypothesis.

Two fMRI datasets were recorded from two cohorts of subjects at two different research sites (University of Florida and University of California, Davis) using the same cued visual-spatial attention paradigm and stimulus parameters. Each trial started with an external cue directing the participants to covertly focus their attention on one of two spatial locations. After a random delay, a stimulus appeared in one of the two locations. The subjects were required to respond to the stimulus if it appeared in the attended location (the attended target) and ignore the stimulus if it appeared in the unattended location (the ignored distractor). Both datasets were preprocessed and analyzed using the same analysis pipeline. Cue-evoked BOLD responses and target-evoked BOLD responses were estimated on a single-trial basis. In the first analysis, MVPA self-decoding and cross-decoding methods were applied to examine pattern decodability within a task period (cue-related task period or target-related task period) as well as pattern cross-decodability between the two task periods (using cue-based classifiers to decode targets or vice versa). The similarity between the cue-evoked and the target-evoked patterns was correlated with behavioral performance to test the attention template hypothesis within visual areas. In the second analysis, the cue-evoked MFC strength was computed and correlated with behavioral performance to examine its functional significance, and the similarity between the cue-evoked MFC pattern and the target-evoked MFC pattern was assessed and linked to behavioral performance to test the attention network template hypothesis.

## Methods

### Overview

Two fMRI datasets were collected at the University of Florida (UF dataset) and at the University of California, Davis (UCD dataset) using the same cued visual spatial attention paradigm (see below). The experimental protocol was approved by the Institutional Review Boards at the respective universities. The participants were neurotypical college students who reported no history of neurological or psychiatric disorders, were right-handed, and had a normal or corrected-to-normal vision. All participants signed written informed consent before taking part in the experiment. They received either course credit or financial compensation for their participation. The UF dataset included thirteen participants (n = 13, 18 to 22 years of age, 5 females and 8 males), and the UCD dataset included eighteen (n = 18, 18 to 22 years of age, 2 females, 16 males). These datasets have been used in previous publications to address different research questions (Bengson et al., 2015; Liu et al., 2016, 2017; Meyyappan et al., 2025; Rajan et al., 2019).

### Experimental paradigm and procedure

The stimuli were shown on an MR-compatible monitor placed outside the scanner bore and viewed via a mirror mounted on the head-coil. Participants were required to fixate a white dot in the center of the screen. Two additional small dots were placed in the lower left and right peripheral visual fields to mark the location where the covert visual spatial attention would be directed to and where the stimulus would appear.

As illustrated in Fig. 1A, each trial started with one of three symbolic cues (circle, square or triangle) displayed slightly above the fixation for 200 ms. Two of the cues, called instructional cues, explicitly instructed the participants to direct their attention covertly either to the left or right visual field location (instructed attention). The third cue, called the choice cue, directed the participants to choose the side of the visual field to covertly attend on that trial (willed attention). The three cue conditions occurred with equal probability, and the meaning of the three symbols was counterbalanced across subjects. Following a stimulus onset asynchrony (SOA) ranging from 2000-8000 ms between cues and target, a vertical grating pattern appeared at one of the two spatial locations marked by the white dot with equal probability for 100 ms. The participants were asked to discriminate the spatial frequency of the target grating (high frequency vs. low frequency) at the attended location via a button press and ignore the grating if it appeared at the unattended location. After the target stimulus by an inter-stimulus interval (ISI) ranging from 2000-8000 ms, participants were prompted to report the location they attended on that trial by the appearance on the screen of “?SIDE?”. The “?SIDE?” prompt permitted a confirmation of the side attended when chosen by the subject, but was also present for the explicitly cued trials for trial consistency. The intertrial interval (ITI) between two trials was also varied randomly across a range from 2000-8000 ms. For the present analysis, the two task periods of interest were the cue period, during which attention control was engaged and active, and the subsequent target period, during which sensory processing and stimulus selection occurred.

**Figure 1.**
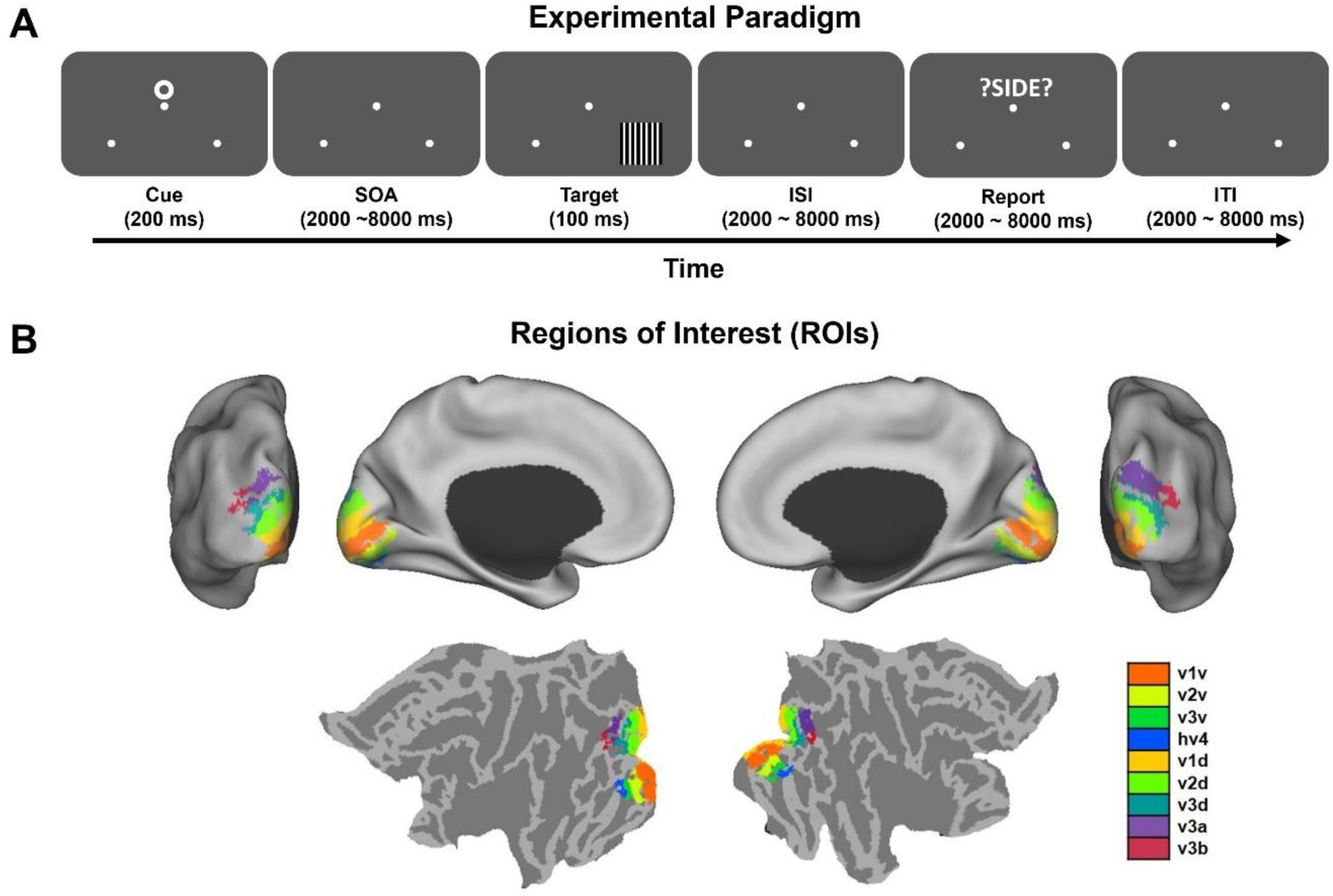
Experimental paradigm and regions of interest (ROIs). (A) Schematic of the trial-by-trial cueing paradigm. Each trial started with a symbolic cue (200 ms) that either directed the subject to covertly attend to the left or the right visual field or directed the subject to choose one of the two visual fields to attend. Following a variable stimulus onset asynchrony (SOA) of 2000-8000 ms, a high-contrast grating was flashed in either the left or right visual field (100 ms). Participants were asked to report the spatial frequency of the grating appearing on the attended side, and ignore the grating on the unattended side. After a second variable interstimulus interval (ISI), a ‘?SIDE?’ prompt was presented and participants were asked to report the visual field they had attended on that trial. The intertrial interval (ITI) varied randomly between 2000-8000 ms. (B) Retinotopic visual cortex, consisting of 9 early visual areas, visualized on the flattened brain. These visual areas were chosen as regions of interests (ROIs).

Prior to fMRI recording, all participants received a training session to familiarize them with the task and to reach a criterion performance of 70% accuracy. The experiment was divided into multiple blocks with rest periods between blocks to prevent participant fatigue. According to the experimental design, there were twice as many instructional trials as choice trials. Because MVPA analyses are more reliable with larger datasets, we focused on the instructional trials. The choice trials were, however, also analyzed, and the results are presented in the Supplementary Materials (Table S1 and Figure S1) in order to provide additional support of the main findings. In addition, by examining the two covert attention conditions (attend left and attend right), which were prompted by the same choice cue, we can eliminate the possibility that the above-chance decoding of cue-related activity in the instructional trials is driven by the difference in the physical appearance of the cues (Carlson & Wardle, 2015).

### FMRI data acquisition and preprocessing

#### UF dataset

FMRI data was recorded on a 3T Philips Achieva scanner (Philips Medical Systems, Netherlands) with a 32-channel head coil using a single-shot echo-planar imaging (EPI) sequence with the following parameters: repetition time = 1980 ms; echo time = 30 ms; flip angle = 80°; field of view = 224 mm; the number of axial slices = 36; voxel size = 3.5×3.5×3.5 mm; matrix size = 64×64. A T1-weighted structural scan was acquired for each subject.

#### UCD dataset

FMRI data was recorded on a 3T Siemens Skyra scanner (Siemens, Erlangen, Germany) with a 32-channel head coil using a single-shot echo-planar imaging (EPI) sequence that has the following parameters: repetition time = 2100 ms, echo time = 29 ms, flip angle = 90°; field of view = 218 mm; the number of axial slices = 34; voxel size = 3.4×3.4×3.4 mm; matrix size = 64×64. A T1-weighted structural scan was acquired for each subject.

Following data collection, the same SPM-based data preprocessing procedure was applied to both datasets, which included the following steps: slice timing correction using sinc interpolation, head motion realignment, co-registration with the standard EPI template, normalization to the MNI space, resampled into 3×3x3 mm voxels, and spatial smoothing with Gaussian kernel with 8mm full width at half maximum. The functional images were further temporally high-pass filtered with the cutoff frequency set at 1/128 Hz. Global signals were adjusted using the proportional scaling approach (Fox et al., 2009).

### Estimation of single-trial BOLD responses

Multivariate pattern analysis (MVPA) requires single-trial data (Kriegeskorte et al., 2006; Norman et al., 2006). The single trial event-related BOLD responses were estimated by using the beta series regression method (Rissman et al., 2004). The BOLD activity evoked by an event of interest (e.g., cues) was modelled using a separate regressor for each trial, and activity evoked by all other events was accounted for by three additional regressors: the left targets and the right targets were modeled by two regressors, and the targets with incorrect responses were modeled by one regressor. For the estimation of target-related BOLD activations, the following regressors were employed: each target was assigned an individual regressor, two regressors were used to model the cues resulting in the participants paying covert attention to the left visual field and the cues resulting in the participants paying attention to the right visual field, and one regressor represented cues in trials where the responses were incorrect.

### Regions of interest (ROIs)

Visual ROIs were defined according to a probabilistic visual retinotopic atlas (Wang et al. 2015). A total of 9 early visual ROIs in the retinotopic visual cortex were included: V1v, V1d, V2v, V2d, V3v, V3d, V3a, V3b, hV4, as shown in Fig. 1B. These areas were chosen based on two considerations: (1) highly precise stimulus information is encoded in these areas (Park & Serences, 2022) and (2) highly precise cue-related information, as indexed by high decoding accuracy between different covert attention conditions, is found in these areas (Meyyappan et al., 2025). These properties render early visual areas ideal for testing the relation between cue-related and target-related neural activity.

### Multivariate pattern analysis (MVPA)

The Linear Support Vector Machine (SVM) was adopted for MVPA decoding analysis. For cue-related activity, the classification or decoding was between the two covert attention conditions: attend left or attend right. For target-related activity, the decoding was between attended left-targets versus attended right-targets or between unattended left-targets versus unattended right-targets. Since training data and testing data came from the same task period, this type of decoding is also called self-decoding, as opposed to cross-decoding described below. Single-trial voxel patterns for a given ROI were input features for SVM classifiers that perform classification by finding a hyperplane to maximize the margin separating two classes of data. Let *k* be the number of voxels in a ROI. Given training data (***X***_*i*_, *Y*_*i*_), for *i* = 1…N (N is the number of trials), the goal of the classifier is to determine the function *f*(***X***) which maps the data ***X***_*i*_ to the class label *Y*_*i*_ such that

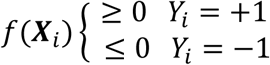

That is, for correct classification, *Y*_*i*_*f*(***X***_*i*_) > 0. For linear SVM, the hyperplane has the form *f*(***X***) = ***w***^*T*^***X*** + *b*, where ***w*** is a *k* dimensional vector normal to the hyperplane and *b* is the bias of the model. From the MVPA classifier we can derive two quantities: decoding accuracy and a weight map. Below we describe how we applied each to test our hypothesis.

#### Decoding accuracy

For each subject, the cue-related data and target-related data were first normalized across trials (subtracting trial-wise mean and divided by trial-wise standard deviation) and then normalized across voxels (subtracting voxel-wise mean and divided by voxel-wise standard deviation) within a given ROI. Normalization across trials is equivalent to mean pattern subtraction which is sometimes called “cocktail-blank removal” (Walther et al., 2016). Mean pattern subtraction moves the center of the data distribution to the origin in the feature space which makes the data from different cognitive conditions (e.g., the cue period and the target period) more comparable. This is important for the cross-decoding analysis in this work (see below). Normalization across voxels removed mean activation to emphasize pattern information. For cue-related decoding analysis, to test whether different attention conditions, e.g., attend left vs. attend right, evoked distinct patterns of neural activity, an SVM classifier was trained and tested on cue-evoked data. Specifically, a 10-fold cross validation approach was used, in which the SVM classifier was trained on nine folds of data and applied to the remaining one-fold of data from the same task period. This process was repeated 10 times using 10 different subsets of trials as testing data, and the decoding accuracies were averaged across the cross-validation folds. To ensure the stability of the decoding results, we repeated 10-fold partition 50 times and averaged all the decoding accuracies to yield the reported decoding accuracy; an above chance decoding accuracy is taken to indicate the presence of the signal of interest (chance decoding accuracy = 50%). For target-related decoding analysis, to test whether the stimulus in the left and the right visual field evoked distinct patterns of neural activity, a similar approach was applied. In what follows, we refer to this type of decoding as self-decoding as training and testing data came from the same task period, to distinguish it from the cross-decoding approach in which the classifier was trained on the data from one task period (e.g., the cue period) and tested on the data from the other task period (e.g., the target period).

Our hypothesis posits that cue-evoked neural patterns are similar to target-evoked neural patterns, and the higher the similarity, the better the match between the cue and the target patterns, leading to better behavioral performance. To examine whether the cues evoked similar patterns as the attended targets, a cross-decoding analysis was applied. For cue-to-target cross-decoding, the training set included all cue-related BOLD responses, and the testing set included all target-related BOLD responses. Because the training and testing sets were from two different task periods, overfitting is not a concern, and therefore cross validation was not necessary to obtain the classification performance. Cross-decoding can also be done by training the classifiers on the target data and testing them on the cue data. An above-chance cross-decoding accuracy is taken to signify similarity between cue-evoked and target-evoked patterns, with higher cross-decoding accuracy indicating higher degree of similarity or pattern matching (chance cross-decoding accuracy=50%).

#### Weight map

To gain more interpretable insights into voxel patterns and their similarity, we also carried out a weight map analysis. As explained above, for a given ROI, the single-trial BOLD activities across voxels are treated as input features for the SVM classifiers, and in this feature space, the weight vector represents the direction orthogonal to the hyperplane pointing in the direction of the positive class. The weight associated with each feature (i.e., a voxel) gives information about the contribution of the voxel to the discrimination of the two attention conditions being decoded. However, the weight vector directly from the SVM is difficult to interpret functionally. A previous study has shown that even when a voxel contains no stimulus information, noise may give the voxel a large weight in the weight vector, because it helps to cancel out the same noise in other voxels which contain stimulus information (Kriegeskorte & Douglas, 2019); see Supplementary Materials for an illustrative example. To make the voxel-level weights more interpretable in terms of the underlying generative cognitive processes, a transformation was applied (Haufe et al., 2014). The corrected weight vector visualized in the form of a brain map is referred to as the weight map henceforth (Lee et al., 2010; Mourão-Miranda et al., 2005). For the two experimental conditions being classified (attend left vs attend right), two populations of voxels can be identified in the weight map: one population is selective for Experimental Condition 1 (e.g., attend left) and the other population for Experimental Condition 2 (e.g., attend right). By being selective for Condition 1, we mean that the BOLD activation in these voxels is higher for Condition 1 than Condition 2, and vice versa. See Figure 2A. The weight map is therefore an intuitive way to visualize the functional brain microstructures underlying the cognitive operation being studied (Rajan et al., 2021). The similarity between the weight maps from cue decoding and target decoding is quantified by cross correlation between the two patterns. A higher cross correlation indicates a higher degree of similarity or better pattern matching in the spatial distribution of the cue-evoked activity and the target-evoked activity.

**Figure 2.**
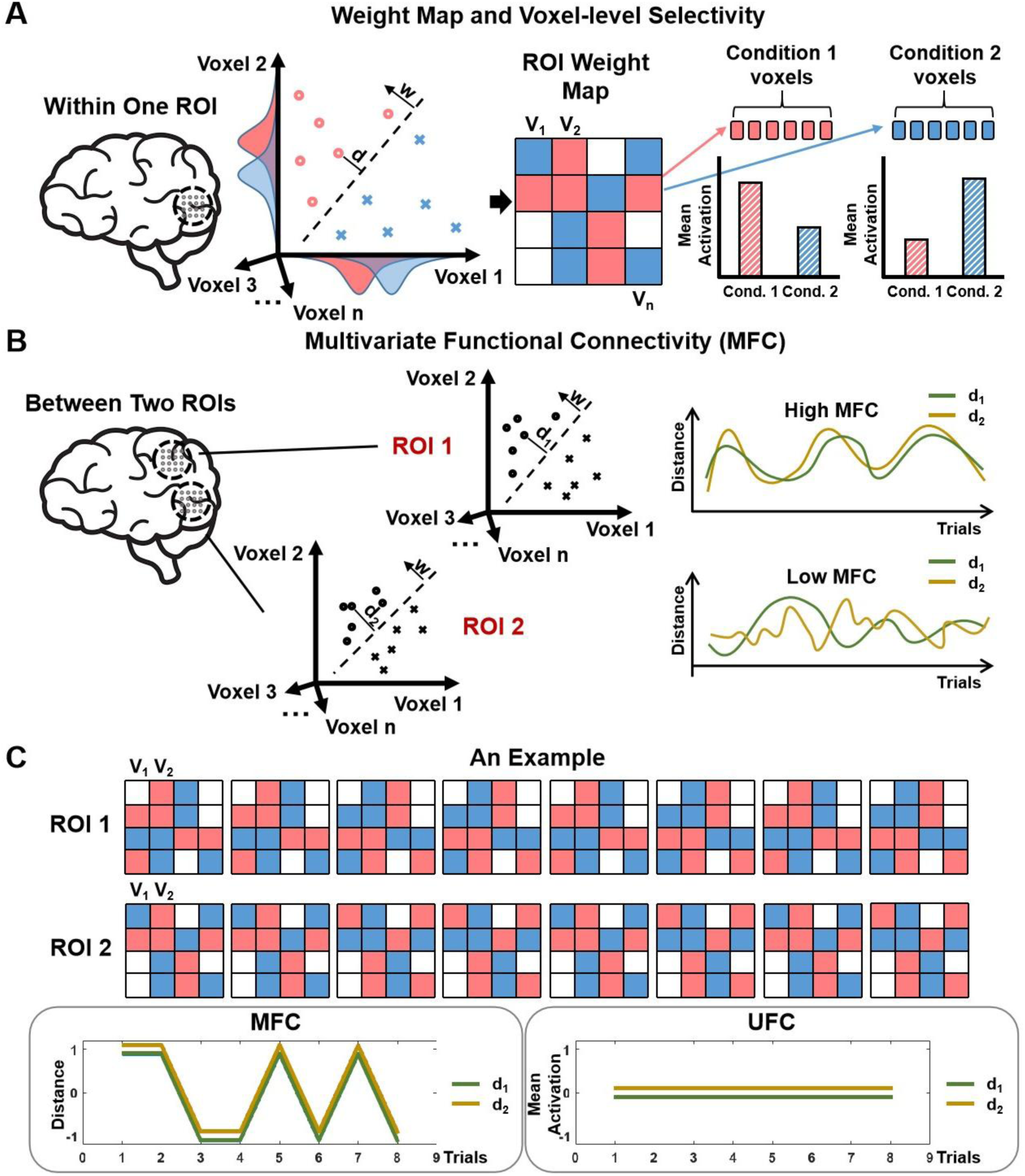
SVM decoding, weight map and functional connectivity. (A) In addition to decoding accuracy, a weight map can be derived from the SVM classifier. Two types of voxels in the weight map identify the two subpopulations of neurons with one population being selective for one of the two experimental conditions and the other being selective for the other experimental condition. (B) Decoding confidence, d_1_ and _2_, from two ROIs, were obtained on a trial-by-trial basis and correlated over trials, yielding multivariate functional connectivity (MFC). A high MFC between two ROIs reflects a higher degree of coordinated pattern activity whereas a low MFC indicates that the two ROIs show weak or no coordinated pattern activity. (C) A simple example where there is strong functional coordination between two ROIs at the multivoxel level (i.e., high MFC) but where the univariate functional connectivity is 0 because the mean activation in each ROI is uniformly zero across trials.

### Functional connectivity analysis

#### Multivariate functional connectivity (MFC)

See Figure 2B. To measure the coordinated activities between two multivoxel patterns from two different ROIs, we computed multivariate functional connectivity (MFC), which had the following steps: (1) Train a SVM classifier in each ROI by using the training data in the cross-validation folds, (2) calculate the signed distance between the test data point to the hyperplane in the feature space for a given trial in the test dataset and denote this signed distance as *d*_1_ for ROI 1 and *d*_2_ for ROI 2, (3) repeat Step 1 & Step 2 for all trials in the test dataset, (4) compute the Pearson correlation between *d*_1_ and *d*_2_ across trials within the testing data fold, (5) average the correlation across all test data folds to yield the MFC between the two ROIs, and (6) transform MFC into z-scores by using the Fisher transformation for statistical comparison and analysis.

The signed distance between the test data point and the hyperplane plane is at the core of MFC, where the sign reflects the side of the hyperplane a test data point falls on, and the distance is sometimes referred to as decoding confidence, because when the sign of the distance aligns with the label of that trial, the larger the distance, the more confident we are about the trial’s class. When *d*_1_ from ROI 1 and *d*_2_ from ROI 2 vary more similarly across trials, meaning that the two ROIs are more coordinated in their pattern level activities, the MFC is larger. Otherwise the MFC is smaller. See Figure 2B for a schematic illustration for two examples with one having high MFC and the other low MFC.

#### Univariate functional connectivity (UFC)

For comparison, we also computed conventional functional connectivity between two ROIs, referred to as univariate functional connectivity (UFC) here. Consider two ROIs. For each trial, the mean activation across voxels within each ROI, denoted as *m*, was computed. The Pearson correlation between the mean activation values from ROI 1 and ROI 2, *m_1_* and *m_2_*, was calculated across trials within a testing data fold and then averaged across all cross-validation folds. Fisher-transformation was performed to enable statistical comparison and analysis. From the definition it is clear that UFC measures the trial-by-trial covariation between the mean activation levels of two ROIs and does not account for pattern-level information within each of the two ROIs.

#### MFC vs UFC

Figure 2C presents a simple example, in which the voxel pattern in each ROI is such that the mean activation is uniformly 0 for both ROIs for all trials, resulting in 0 UFC. However, there is strong functional coordination between the ROIs at the neural pattern level, which is revealed by MFC.

### Combining datasets using meta-analysis

We analyzed two datasets recorded at two different institutions to (1) demonstrate reproducibility of the findings and (2) increase statistical power when needed to generate more statistically reliable findings. For the latter we combined the two datasets via meta-analysis by using the Lipták-Stouffer method (Lipták, 1958). The method involves converting the p-value from each dataset to its corresponding z value and combining the z values from the two datasets using the Liptak-Stouffer formula:

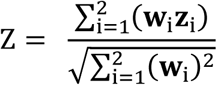

Here Z represents the aggregate z-score from the two datasets, **z**_i_ refers to the z value from the *i*th dataset (i=1, 2), and **w**_i_ is the weight from the *i*th dataset which is equal to the square root of the sample size in the *i*th dataset. From the Z score, the meta p value is assessed, and the statistical significance of the result determined.

## Results

Subjects were cued to covertly attend the left or the right visual field (instructed trials) or asked to choose to attend one of the two visual fields (choice trials); cue-left trials, cue-right trials, and choice trials were equally likely. After a random delay, a stimulus appeared in one of the two visual fields, and the subjects discriminated the stimulus appearing in the attended visual field (attended target) and ignored the stimulus appearing in the unattended visual field (ignored or unattended target). Because there were twice as many instructed trials as choice trials, to maximize our statistical ability to test the hypotheses, the instructed trials were the main focus here. Choice trials were also analyzed and presented in the Supplementary Materials to provide further support for the main conclusions.

### Behavioral analysis

**Table 1.** Behavioral results.

| Dataset | Condition | Reaction Time (ms) | Accuracy (%) |
| --- | --- | --- | --- |
| UF | Cue (Attend) Left | 918.56±24.49 | 85.72±2.14 |
|  | Cue (Attend) Right | 911.81±23.28 | 85.80±1.24 |
| UCD | Cue (Attend) Left | 1027.02±55.60 | 82.45±2.30 |
|  | Cue (Attend) Right | 1041.87±54.95 | 81.97±2.10 |

#### UF dataset

For cue-left and cue-right trials, the target discrimination accuracy was 85.72 ± 2.14% and 85.80 ± 1.24% and the reaction time was 918.56 ± 24.49 ms and 911.81 ± 23.28 ms, respectively. There was no statistically significant difference between cue-left and cue-right trials in both target discrimination accuracy and reaction time (p_ACC_ = 0.97, p_RT_ = 0.67).

#### UCD dataset

For cue-left and cue-right trials, the target discrimination accuracy was 82.45 ± 2.30% and 81.97 ± 2.10% and the reaction time was 1027.02 ± 55.60 ms and 1041.87 ± 54.95 ms, respectively. There was no statistically significant difference between cue-left and cue-right trials in both target discrimination accuracy and reaction time (p_ACC_ = 0.67, p_RT_ = 0.50).

### Self-decoding of cue-related and target-related BOLD activities

To test whether covert attention to the left visual field and the right visual field generated distinct neural patterns in the visual cortex, we employed a self-decoding approach in which we trained and tested classifiers using data from the cue period; a 10-fold cross validation was applied to minimize overfitting. As shown in Figure 3A, across the retinotopic visual areas tested, the decoding accuracy between cue-left vs cue-right was well above chance level of 50%, suggesting that the top-down biasing signals were present in these visual areas before target onset, and the results are consistent across the two datasets, indicating a high degree of reproducibility. It is worth noting that cue decoding has been presented in a previous publication (Meyyappan et al., 2025), it is included here for completeness, and helps to facilitate the comparison between cue-related and target-related activities, which is the main purpose of this study.

**Figure 3.**
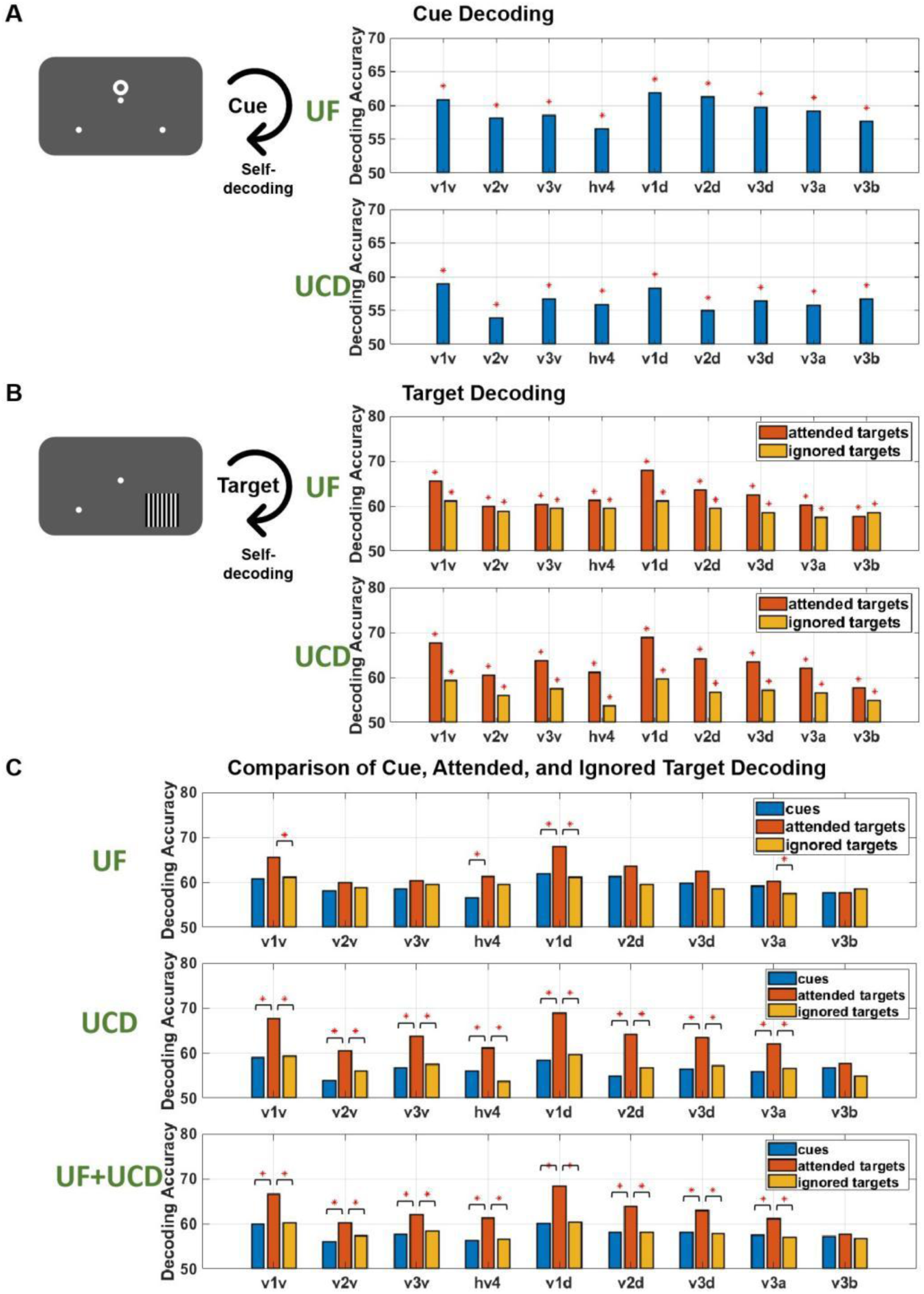
Self-decoding of cue-related and target-related neural activities. (A) Accuracy of decoding attend-left vs attend-right in the cue period. The direction of covert attention can be decoded in all ROIs. The results are consistent across the two datasets. (B) Accuracy of decoding attended left-target vs attended right-target and ignored left-target vs ignored right-target in the target period. The location of both the attended target and the unattended target can be decoded in all visual ROIs. The results are consistent across the two datasets. (C) Comparison of cue decoding accuracy, attended target decoding accuracy, and ignored target decoding accuracy. Combining the two datasets via meta-analysis, attended target decoding accuracy was significantly higher than cue decoding accuracy and ignored target decoding accuracy in all ROIs except v3b where no significant difference was found. Self-decoding here means that the classifiers were trained and tested on the data from the same task period. *: p<0.05.

To test whether the target appearing in the left visual field and in the right visual field generated distinct neural patterns and whether attention modulated these patterns, we applied a similar self-decoding approach to BOLD activity patterns evoked by the attended targets and by the ignored targets. The results showed that the decoding accuracy between the attended left-target vs attended right-target as well as between the ignored left-target vs ignored right-target was significantly above chance (Figure 3B), suggesting that the target stimulus appearing in different spatial locations evoked distinct neural patterns, irrespective of whether it is attended or not. In Figure 3C, a meta-analysis combining the two datasets demonstrated that the attended target decoding accuracy is significantly higher than that of the ignored target in all visual ROIs except v3b where no significance difference was found, suggesting that attention enhanced differences in neural patterns evoked by the target in the two different visual fields. In addition, as shown in Figure 3C, the decoding accuracy for the attended targets was significantly higher than that for the cues for all ROIs except v3b where no significant difference was found, demonstrating that the attended target appearing in two spatial locations evoked more strongly distinct neural patterns than deploying anticipatory covert attention to the same two spatial locations.

### Cross-decoding between cue-related and target-related BOLD activities

Our attention template hypothesis posits that cue-evoked activity patterns are similar to the activity patterns evoked by the attended target. This was tested by conducting a cross-decoding analysis. Classifiers were trained on the attended targets (attended left-target vs attended right-target) and then applied to decode the cues (cue-left vs cue-right). As shown in Figure 4A, the cross-decoding accuracies are significantly above chance level in all ROIs. Similar observations were obtained by applying cue-trained classifiers to cross-decode attended targets (Figure 4B). Importantly, classifiers trained on cues did not decode ignored targets, nor did classifiers trained on ignored targets decode cues, suggesting that the neural patterns evoked by the ignored target are different from that evoked by the attention-directing cues. Notably, these results are consistent across the two datasets.

**Figure 4.**
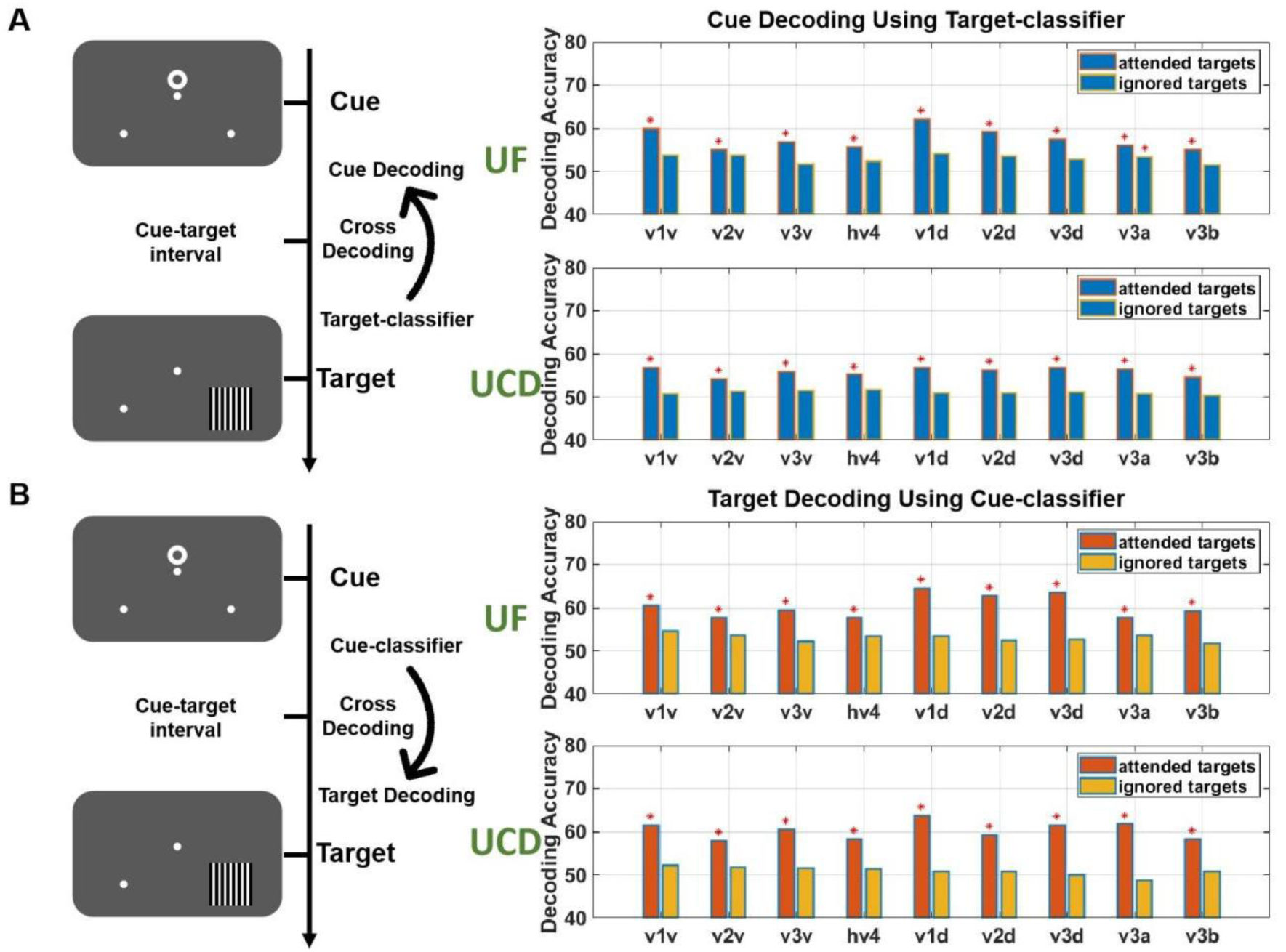
Cross-decoding between cue-related and target-related neural activities. (A) Classifiers trained on attended targets decode cues in all ROIs. Classifiers trained on ignored targets, on the other hand, do not decode cues. The results are consistent across the two datasets. (B) Classifiers trained on cues decode attended targets but not ignored targets in all ROIs. The results are consistent across the two datasets. *: p<0.05.

### Weight map analysis

Although above-chance cross-decoding accuracy implies similarity between neural patterns evoked by attention-directing cues and the attended visual stimuli, a weight map analysis can shed additional lights on the underlying neural mechanisms. To illustrate, consider the left dorsal V1 (left v1d) as an example ROI. For a typical subject, Figure 5A shows the weight map from the cue-trained classifier (decoding cue-left vs cue-right) and that from the target-trained classifier (decoding attend left-target and attended right-target). The similarity between the two weight maps is apparent and is supported quantitatively by the very high correlation coefficient of R = 0.91. As explained in Figure 2A, from the cue weight map, the voxels within the ROI can be categorized into two groups: cue-left voxels and cue-right voxels. The average BOLD activities were computed within each group of voxels for cue-left trials and cue-right trials. As shown in Figure 5B, larger BOLD signals were evoked by the attending-left cue compared to the attending-right cue in the cue-left voxels, whereas larger BOLD signals were evoked by the attending-right cue compared to the attending-left cue in the cue-right voxels. Similarly, from the target weight map, the voxels within the ROI can be categorized into two groups: left-target voxels and right-target group, and the average BOLD activities were computed within each group of voxels for attended left-target trials and attended right-target trials. Figure 5B shows that larger BOLD signals were evoked by the attended left-target compared to the attended right-target in the left-target voxels, whereas larger BOLD signals were evoked by the attended right-target compared to the attended left-target in the right-target voxels. The similarity between the cue weight map and the target weight map suggest that the voxels selective for the left or the right target are also selectively activated during covert attention to the left or the right visual field, lending further support to our attention template hypothesis. In Figure 5C, we correlated the cue weight map and the attended target weight map in all ROIs for each subject and averaged the results across all subjects, and found significant correlation across ROIs, suggesting that the similarity between cue-evoked patterns and target-evoked patterns is a general property across early visual cortex. Notably, these results are consistent across the two datasets.

**Figure 5.**
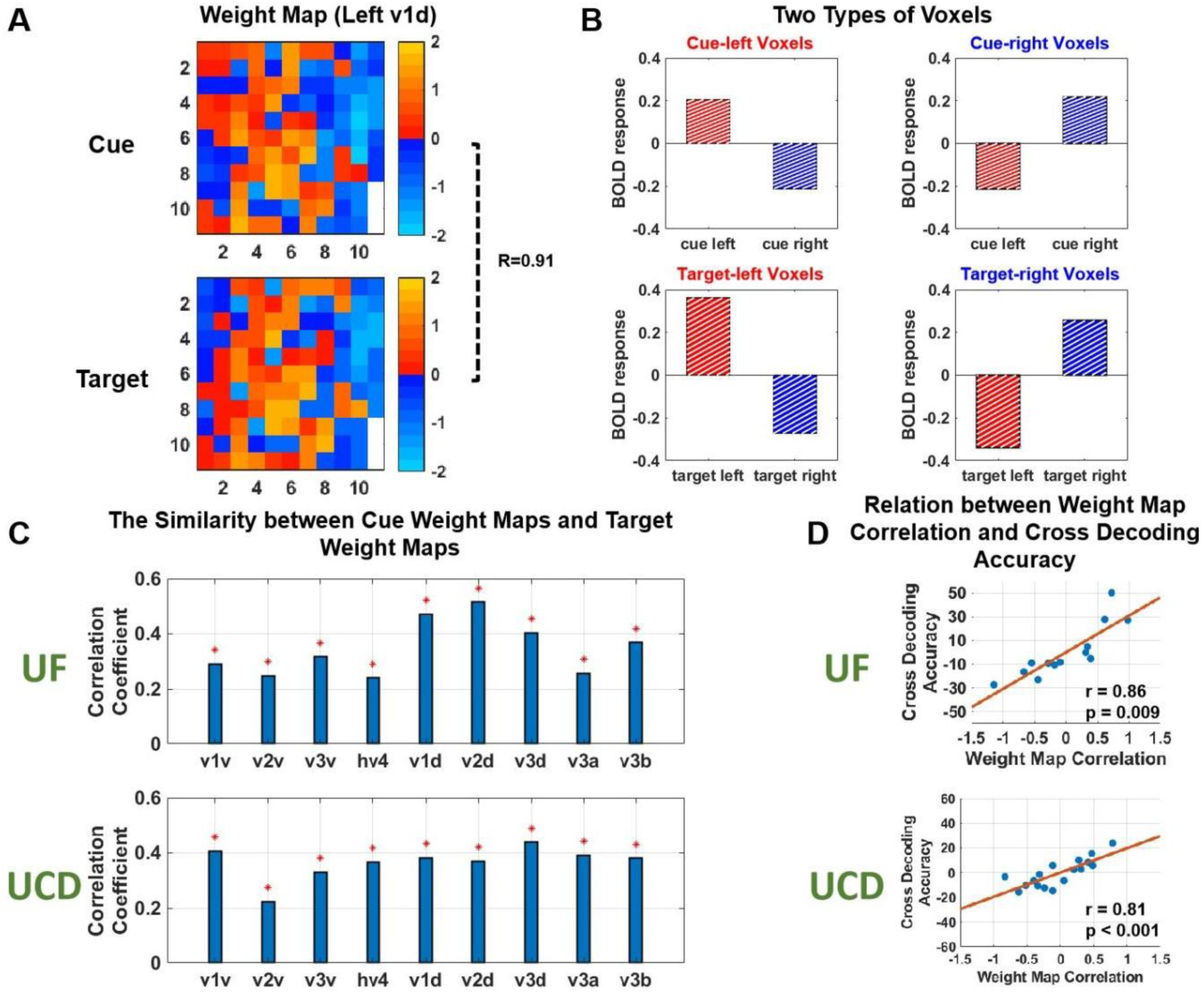
Weight map analysis. (A) Cue weight map and attended target weight map from a typical subject for the ROI of left v1d. The high degree of similarity between the two weight maps is quantified by a correlation of R=0.91. (B) Voxel-level selectivity derived from the weight map. (C) Significant correlation was found at the population level between cue weight map and attended target weight map for all ROIs. (D) The relation between cross-decoding accuracy and weight map correlation quantified by the respective first PCA component correlation. *: p<0.05.

### Weight map correlation vs cross-decoding

We have thus used both cross-decoding and weight map correlation to quantify the similarity between cue-evoked patterns and attended target-evoked patterns. As indicated earlier, these two measures should be related. We tested this in Figure 5D. Because each ROI has a cross-decoding accuracy and a weight map correlation, to perform the analysis at the whole visual cortex level - at the same time to reduce dimension, we note that the weight map correlation and cross-decoding accuracy covary across ROIs, thereby providing the basis for carrying out a PCA analysis on both the weight map correlation and the cross-decoding accuracy. For the UF dataset, the first PCA component of the weight map correlation and the cross-decoding accuracy explained 58.38% and 79.48% of the data variance, whereas for the UCD dataset, the first PCA component of the weight map correlation and the cross-decoding accuracy explained 32.88% and 48.88% of the data variance. As shown in Figure 5D, across subjects, the correlation coefficient between the score on the first weight map correlation PCA component and the score on the first cross-decoding accuracy PCA component was r_UF_ = 0.86 (p = 0.009) for UF dataset and r_UCD_ = 0.81 (p < 0.001) for UCD dataset. The high correlation between the two measures was thus demonstrated for both datasets.

### Cue-target pattern matching and behavior

Our attention template hypothesis further posits that the cue-evoked pattern facilitates the selection of attended information via a pattern matching mechanism, which means that the higher the similarity between the cue-evoked and the attended target-evoked patterns, the more effective the stimulus selection, the better the behavioral performance. To test this, we used the score on the first PCA component of the weight map correlation to index the individual difference in cue-target pattern similarity and correlated it with the reaction time. As shown in Figure 6A, across subjects, a significantly negative correlation was found for both datasets (r_UF_ = −0.56, p_UF_ = 0.046; r_UCD_ = −0.67, p_UCD_ = 0.002), suggesting that the better the cue-target pattern matching in the visual cortex, the faster the reaction time.

**Figure 6.**
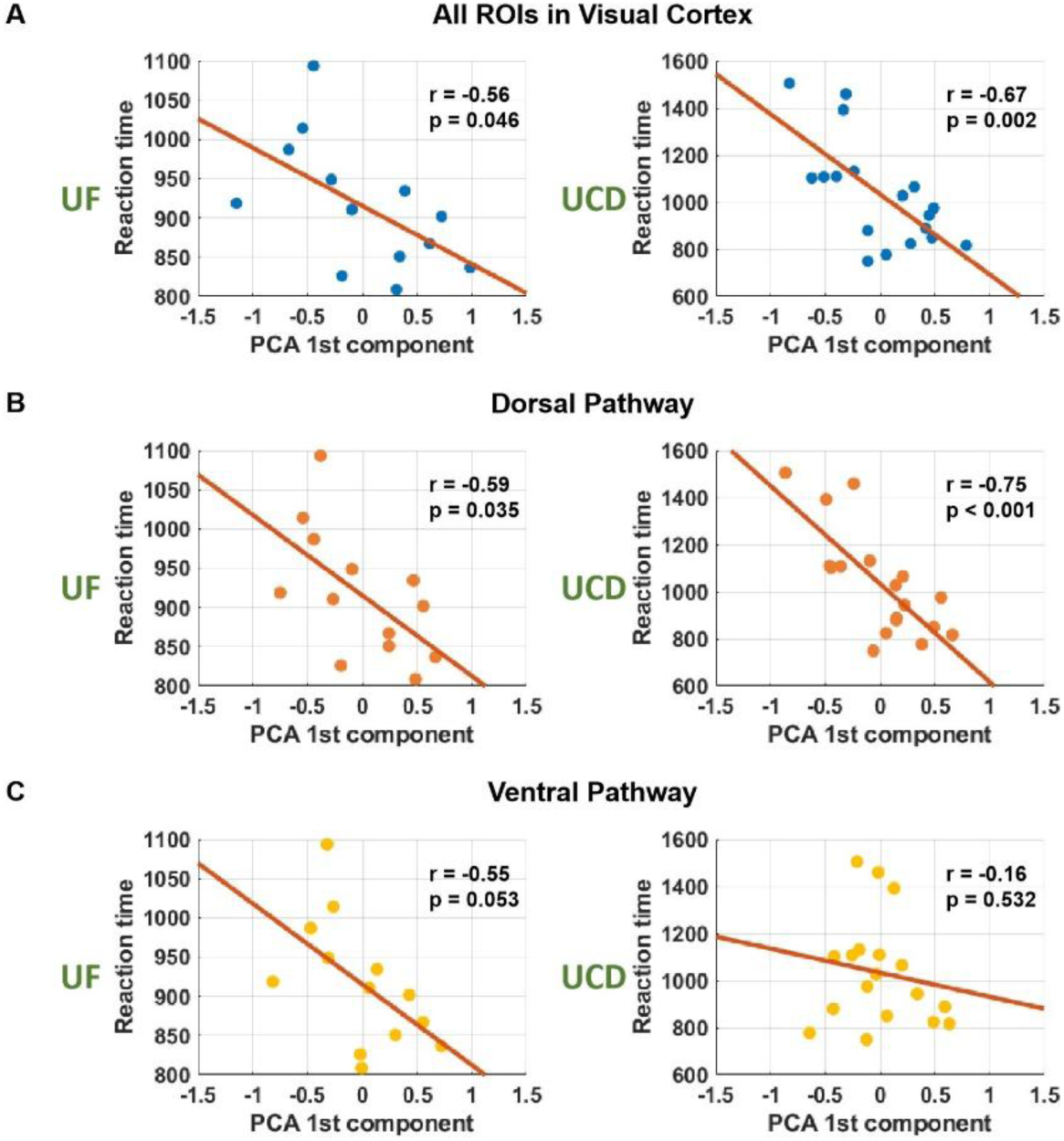
Cue-target pattern similarity (template matching) and behavior. (A) The relation between cue-target pattern matching in visual cortex and reaction time. The PCA analysis was applied to all ROIs and the score on the first PCA component was used to index cue-target pattern similarity or template matching. Significant negative correlation was observed in both datasets (better template matching, faster reaction). (B) The relation between cue-target pattern matching in dorsal pathway and reaction time. Significant negative correlation was observed in both datasets. (C) The relation between cue-target pattern matching in ventral pathway and reaction time. No significant correlation was observed in either dataset.

To further assess the contribution of the ventral vs dorsal pathways to behavioral facilitation, we divided ROIs into dorsal ROIs: v1d, v2d, v3d, v3a, v3b and ventral ROIs: v1v, v2v, v3v, hv4, and performed the PCA analysis within each pathway. For the UF dataset, the first PCA component explained 63% and 62% of the variance within the dorsal and the ventral pathways, whereas for the UCD dataset, the variance explained was 48% and 47%. As shown in Figure 6B, a significantly negative correlation between the cue-target pattern matching in the dorsal pathway and the reaction time was observed for both datasets (r_UF dorsal_ = −0.59, p_UF dorsal_ = 0.035; r_UCD dorsal_ = −0.75, p_UCD dorsal_ = 0.001). In the ventral pathway (Figure 6C), however, the cue-target pattern matching and the reaction time was not significantly correlated for either of the two datasets (r_UF ventral_ = −0.55, p_UF ventral_ = 0.053; r_UCD ventral_ = −0.16, p_UCD ventral_ = 0.532); a meta-analysis combining the two datasets yielded p_ventral_ = 0.189, which again showed no significant correlation. These results suggest that whereas cue-target pattern matching takes place within both dorsal and ventral pathways, it is the cue-target pattern matching within the dorsal pathway that positively contributes to the facilitation of behavioral performance (Mishkin et al., 1983).

### Functional connectivity analysis

To test the attention network template hypothesis, we applied multivariate functional connectivity (MFC) analysis to fMRI data. Since MFC is a relatively new measure, we first applied it to cue-evoked activity, and tested its functional relevance by relating the result to behavior. Given that MFC is based on classifiers trained to decode cue-evoked attend left vs attend right, both types of cues were used to calculate MFC. Conducting a PCA analysis on cue evoked MFC across subjects, the first PCA component explained 83.28% of the data variance for the UF dataset and 62.97% of the data variance for the UCD dataset. As shown in Figure 7, a negative correlation was found between the score on the first PCA component of the cue-evoked MFC and reaction time, where r_UF_ = −0.49, r_UCD_ = −0.67, and the meta-analysis p = 0.001, suggesting that the stronger the pattern level functional connectivity in the cue period, the better the behavioral performance indexed by shorter reaction time. For comparison, cue-evoked univariate functional connectivity (UFC) was computed for all pairs of visual ROIs, and the score on the first PCA component of UFC, which explained 58.34% of the data variance for the UF dataset and 37.26% of the data variance for the UCD dataset, exhibited no correlation with behavior, where r_UF_ = 0.08, r_UCD_ = 0.08, and the meta-analysis p = 0.842. These results demonstrated that it is important to take into account the pattern-level coordinated activity across the visual cortex and the MFC considered here is a good measure of the pattern-level coordinated activity. We then applied MFC to target-evoked activity. As shown in Figure S2 (Supplementary Materials), attention enhanced functional connectivity in 19 out of 36 ROI pairs measured by MFC but 0 ROI pair measured by UFC, further demonstrating the importance of taking into account pattern-level information and the utility of MFC.

**Figure 7.**
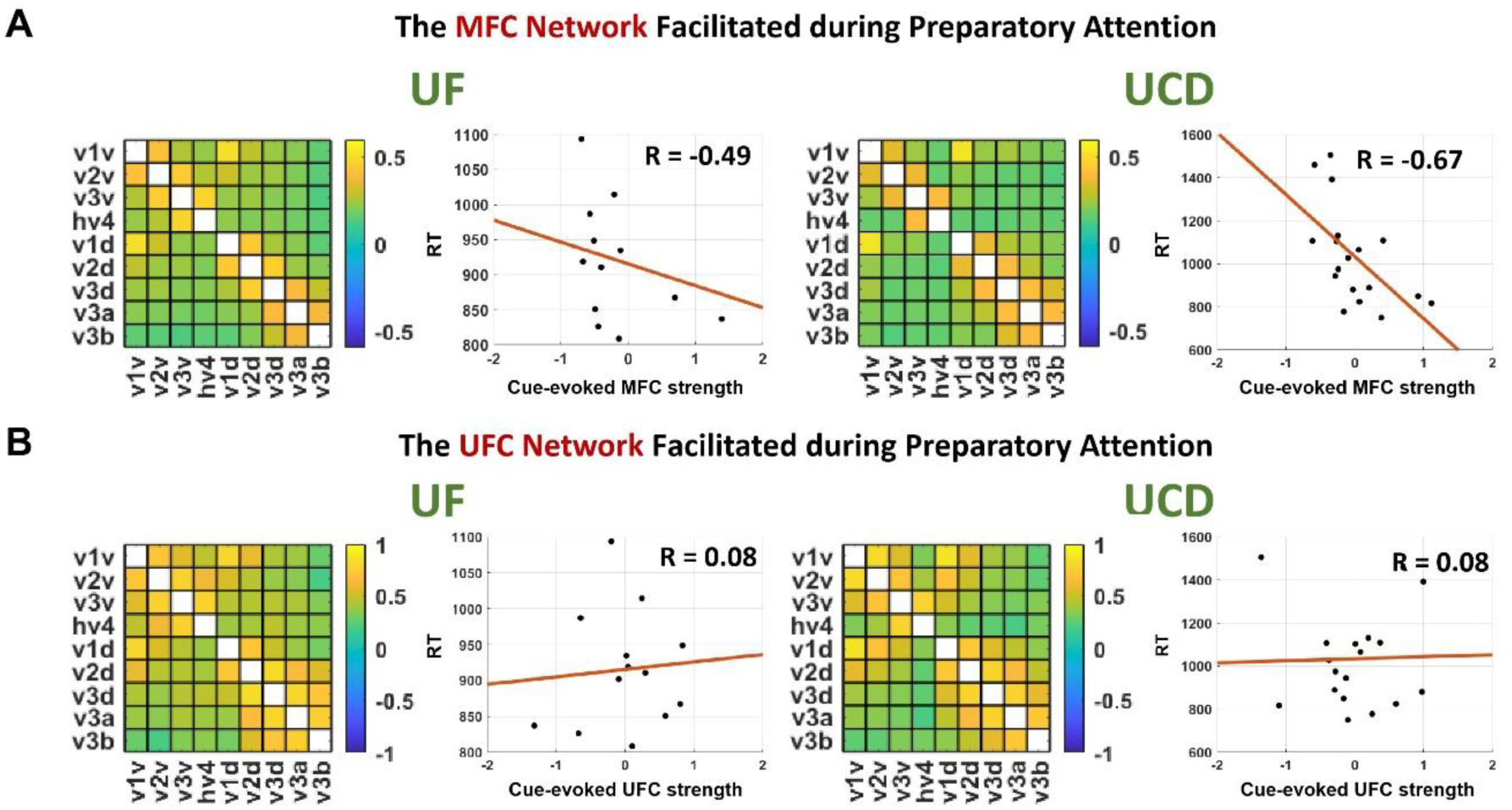
Functional connectivity analysis in the cue period. (A) MFC of all pairs of ROIs in visual cortex evoked by the cues and the relationship between mean cue-evoked MFC strength and reaction time. (B) UFC of all pairs of ROIs in visual cortex evoked by the cues and the relationship between mean cue-evoked UFC strength and reaction time.

Our attention network template hypothesis posits that (1) the cue evokes a functional connectivity pattern that is similar to the functional connectivity pattern evoked by the attended target and (2) the higher the similarity between the functional connectivity patterns from the two task periods, the more effective the network level stimulus selection, the better the behavioral performance. We tested this hypothesis using MFC. For the cue period, MFC was computed for each ROI pair based on the classifiers trained on cue-evoked data to decode attend left vs attend right, and the result for each subject was averaged across subjects. For the target period, MFC was computed for each ROI pair based on the classifiers trained on attended target-evoked data to decode attended left target vs attended right target, and the result for each subject was averaged across subjects. Figure 8A shows cue-evoked MFC and attended target-evoked MFC across ROIs. The strong correlations consistent across the two datasets, r_UF_ = 0.93, r_UCD_ = 0.95, p << 0.05, suggest a high degree of similarity at the population level between the cue-evoked functional connectivity pattern and the attended target-evoked functional connectivity pattern. At the individual subject level, Figure 8B shows the results from two example subjects from the UF dataset, one with high and the other low similarity between cue-evoked and attended target-evoked MFC connectivity patterns. The subject with high cue-target MFC network similarity has better behavioral performance (RT=867 ms) than the subject with low cue-target MFC network similarity match (RT=1014 ms). Across subjects, as shown in Figure 8C, a significant negative correlation was found between cue-target MFC network similarity and reaction time, with r_UF_ = −0.35, r_UCD_ = −0.51, and the meta-analysis p = 0.029, suggesting that when the signal processing pathways are more activated during preparatory attention, there is a greater enhancement in task performance. Our attention network template hypothesis was thus supported.

**Figure 8.**
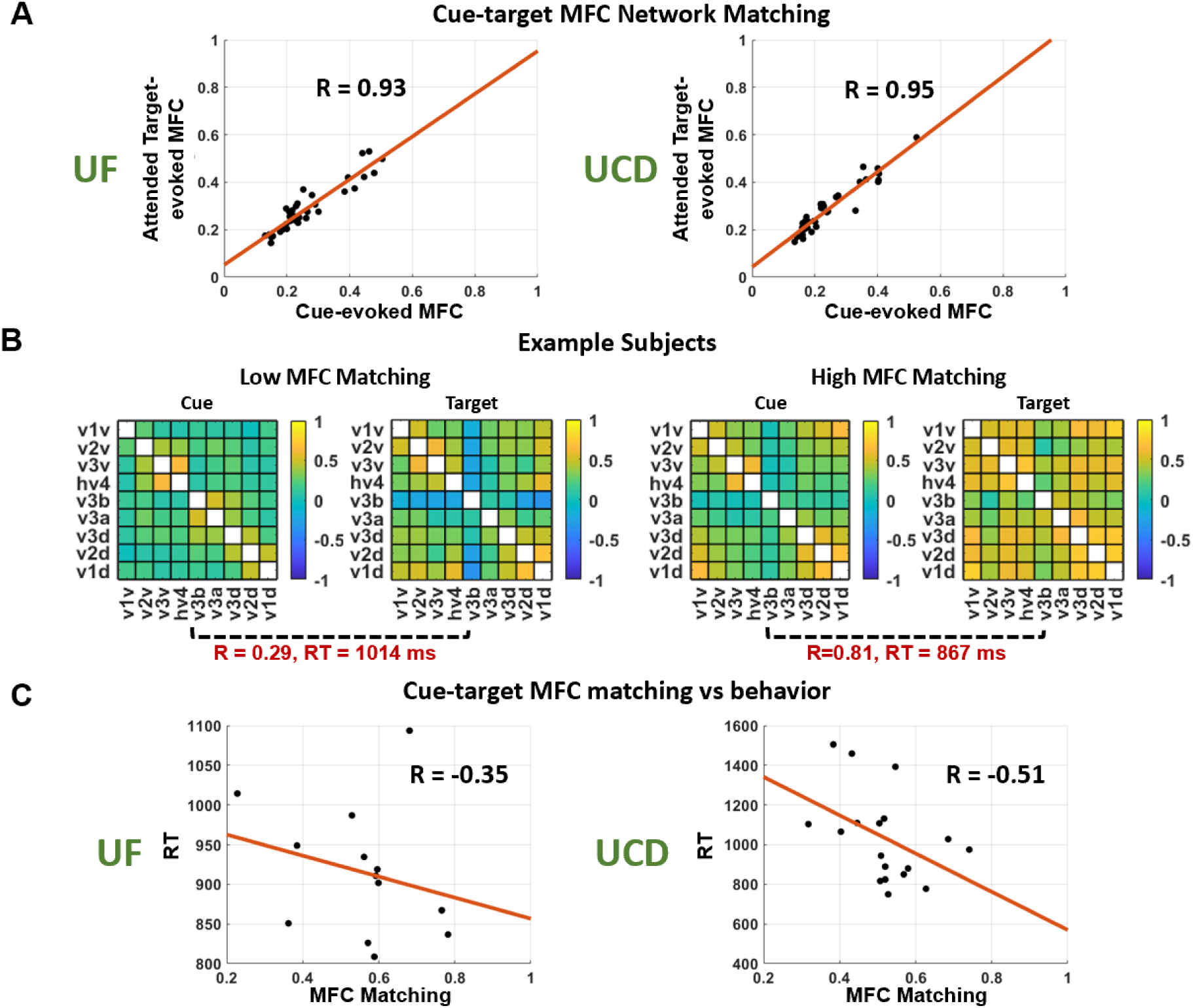
Cue-target functional connectivity pattern similarity and relation with behavior. (A) The relationship at the population level between MFC in the cue period and that in the target period; here each dot represented one ROI pair. (B) Two example subjects showing high versus low MFC network matching and their behavioral performance. (C) Across subjects, a significant negative correlation was found between cue-target MFC network similarity and reaction time; here each dot represented one subject).

For comparison, we performed the same analysis using UFC instead of MFC. The cue-evoked and attended target-evoked UFC patterns were also highly correlated across ROIs at the population level, where r_UF_ = 0.99, r_UCD_ = 0.99, p << 0.05, suggesting strong similarity between the functional connectivity patterns activated during the two different task periods. However, across subjects, no significant correlation was found between cue-target UFC network similarity and reaction time, with r_UF_ = −0.27, r_UCD_ = −0.24, and the meta-analysis p = 0.291.

## Discussion

Voluntary visual-spatial attention cueing paradigms typically consist of two tasks periods: the cue period in which the attention control is active and the target period in which stimulus selection takes place. Most models posit that top-down attention control biases neural activities in visual cortex during the cue period in anticipation of the target stimulus, leading to selective stimulus processing when the target appears. We investigated how attention biasing in the visual cortex facilitated stimulus selection and behavioral performance. Analyzing fMRI data using cross-decoding MVPA methods, we found that in the visual cortex, the patterns of neural biasing activity evoked by the spatial cues directing covert attention to the left or the right visual field were similar to the patterns of neural activity evoked by the attended target appearing in the left or the right visual field. Higher similarity between the neural patterns from the two task periods was correlated with better behavioral performance. Further, the functional connectivity patterns triggered by the cues and by the attended targets in visual cortex were similar across all possible pairs of visual ROIs as quantified by multivariate functional connectivity (MFC). Higher similarity between the functional connectivity patterns from the two task periods also correlated with better behavioral performance.

### From sensory biasing to stimulus selection

When attention is deployed in advance of sensory stimulation, as in typical cue-target paradigms, the subsequent stimulus processing is enhanced and behavioral performance improved (Carrasco, 2011). Neurophysiologically, according to the prevailing model, following an attention directing cue, top-down signals propagate from the DAN to the visual cortex to bias sensory processing in favor of the attended information (Battistoni et al., 2017; S. L. Bressler et al., 2008; Carrasco, 2011; Chawla et al., 1999; Giesbrecht et al., 2006; Hopfinger et al., 2000; Kastner et al., 1999; Luck et al., 1997; Munneke et al., 2008; Park & Serences, 2022; Sylvester et al., 2007; Ungerleider, 2000). In support of the theory, evidence from single-cell studies showed that spatial attention increased the firing rate of neurons in V2 and V4 even in the absence of stimulus input (Connor et al., 1997; Luck et al., 1997). In humans, Hopfinger et al. (2000) showed that when different attention-directing cues are contrasted, BOLD activity in higher-order, but not lower-order, visual areas are elevated by attention. In lower-order visual areas such as V1, it has been reported that directing covert attention to a spatial location increased BOLD activity in some voxels but decreased activity in other voxels, suggesting that cue-related attention modulation may be more fruitfully studied at the neural pattern level (Müller & Kleinschmidt, 2004; Silver et al., 2007; Smith et al., 2000). Applying MVPA techniques to fMRI data from the cue period of the same datasets analyzed here, we recently reported that cue-related biasing represented as changes in BOLD signal patterns can be decoded in all retinotopic visual areas, and lower-order visual areas actually have higher decoding accuracy than higher-order areas, suggesting that (1) attention biasing is prevalent throughout the visual hierarchy, and (2) the strength of attention biasing, as indexed by decoding accuracy, may follow a reverse hierarchy pattern (Meyyappan et al., 2025).

What is the relation between sensory biasing during anticipatory attention and subsequent stimulus processing? William James likened anticipatory attention to visual features or objects to an “image in the mind” (James, 2007). This mental image, sometimes referred to as an attention template, is thought to be implemented by the selective activation of neurons within the visual cortex prior to the processing of sensory information. Using MVPA decoding methods, it has been shown that for stimuli as wide ranging as letters, shapes or objects, neural patterns following the deployment of attention but before the appearance of the target closely mirror the brain activity patterns observed during the perception of the same stimuli (Gayet & Peelen, 2022; Peelen & Kastner, 2011; Stokes et al., 2009), lending support to the attention template idea; but see Gong et al. (2022) for evidence supporting the alternative idea that these activity patterns are non-sensory in nature(Gong et al., 2022). The present work addressed the question of whether the attention template idea is applicable to visual spatial attention. Our results appear to answer the question in the affirmative. Applying the MVPA approach to fMRI data, we showed that neural patterns evoked by cues directing attention to different spatial locations resemble that evoked by the attended target stimuli appearing at those locations, both in terms of above-chance cross-decoding accuracy and significant weight map correlation. In addition, treating the magnitude of the cue-target weight map correlation as a measure of the degree of pattern matching between the cue-evoked pattern and the target-evoked pattern and the reaction time as a measure of behavioral performance, we further showed that there was a negative correlation between the two, suggesting that stimulus selection is by template matching, namely, the better the stimulus pattern matches that activated by the cue, the faster the reaction time. We note that these results can also be seen as substantiating a recent suggestion by Jimenez et al. (2024) that there exist location templates in visual search and these location templates facilitate behavior equally well as the more classic feature-based color templates.

### From anticipatory network facilitation to stimulus selection

There are at least two non-mutually exclusive views on how attention improves perception. Focusing on an individual visual area, the conventional view posits that attention enhances perception by prioritizing certain information processed by the area, e.g., by amplifying the evoked response that represents the information (Carrasco, 2011; Mangun & Hillyard, 1991; McAdams & Maunsell, 1999; Saenz et al., 2002; Spitzer et al., 1988; Treue & Maunsell, 1996; Van Voorhis & Hillyard, 1977). Another view suggests that attention modifies communications between brain areas (Womelsdorf & Fries, 2007). This view stems from the observation that attention is linked to increased synchrony between sensory areas and between sensory and higher-level cortical areas. For example, studies have demonstrated that spatial attention improves the efficacy of thalamo-cortical sensory transmission (Briggs et al., 2013) and modulates synchrony between V1 and V4 (Bosman et al., 2012) and that attention to motion boosts synchrony between V1 and MT through spike count correlation (Ruff & Cohen, 2016)

Our attention network template hypothesis seeks to address the relation between the functional connectivity pattern during attention control (cue period) and that during stimulus selection (target period). In line with models of network-level attention processes based on synchrony (Fries, 2015), it posits that top-down attention control generates sensory biasing signals that pre-activate the transmission pathways of the attended target stimulus, and that this takes the form of connectivity patterns (network templates) of neural facilitation and suppression of connections among visual areas. The closer the pre-activated network connectivity pattern resembles the target-evoked connectivity pattern, the more efficiently stimulus-related information flows within visual areas and from sensory processing to motor execution, resulting in better behavioral performance (e.g., faster reaction times). To test this, we first noted that the traditional functional connectivity measure (i.e., ROI-level cross correlation) ignores pattern-level information, and neural patterns form the basis of our attention template hypothesis. Consequently, a multivariate functional connectivity measure was defined and applied, which measures coordinated activities between visual areas at the neural pattern level. By correlating the cue-evoked MFC functional connectivity with the reaction time across subjects, we found that those with a stronger preactivated signaling network before stimulus onset responded to the stimulus more quickly. More importantly, the detailed connectivity pattern of the preactivated network closely resembles that of the stimulus processing network evoked by the attended target, and the similarity between the network connectivity patterns from the two task periods is negatively correlated with reaction time, i.e., the better the cue-evoked network pattern matches the target-evoked network pattern, the faster the reaction time. These results are in support of the attention network template idea and demonstrate a significant role of the pre-activation of the signal transmission pathway during preparatory attention in stimulus selection.

### Additional findings

Two additional findings on attentional modulation of stimulus processing are worth noting. First, we found that unattended targets also evoked decodable patterns of neural activity, but the decoding accuracy is significantly lower than for attended targets. The stronger pattern differences between the attended left-target and the attended right-target reflect that the voxels selective for a given target (e.g., left-target) are further enhanced by attention to that target whereas the voxels not selective for that target are further suppressed by attention to that target. This agrees well with Bressler et al. (2013) who found that spatial attention enhances both positive and negative BOLD signals to the attended relative to the unattended visual stimulus (Bressler et al., 2013). Second, we found that attended targets evoked significantly higher multivariate functional connectivity among visual ROIs than ignored targets; that is, attention enhanced MFC among 19 out 36 total possible pairwise ROI connections. For univariate functional connectivity (UFC) analysis, no attention modulation was found, suggesting that attention increased pattern level functional coordination in visual cortex rather than mean ROI level functional coordination (Figure S2). This fact further highlights the importance of considering neural activity at the pattern level when performing functional network analysis.

### Methodological considerations

First, although cross-decoding is an established method for assessing pattern similarity across two different task periods, the weight map provides a way to visualize the voxel patterns from both task periods and gives functional significance to the voxels in the ROI. In particular, the Haufe et al. transformation (2013) allowed us to divide the voxels into two groups, with each group of voxels being selectivity for one of the two experimental conditions being decoded. Based on this method, we are able to suggest that the voxels selective for an attended stimulus are also preactivated during covert attention to the same spatial location, whereas the voxels not selective for the given attended stimulus are suppressed at the same time. This extends single unit studies (Chelazzi et al. 1993, 1998) showing that neurons in the inferior temporal cortex selective for a stimulus exhibit higher baseline activity during the preparatory attention when cued to attend to that stimulus. Second, recognizing that the univariate functional connectivity ignores the multivoxel pattern information, previous work has proposed various multivariate connectivity methods (Anzellotti & Coutanche, 2018; Basti et al., 2020; Fang et al., 2023). The MFC method formulated in this work is one such method that addresses the limitation of the traditional univariate functional connectivity method. Having demonstrated that coordinated brain activity in visual cortex during the cue period is functionally significant, we further showed that MFC played a crucial role in testing the attention network template idea by showing that pattern matching between cue-evoked MFC connectivity pattern and target-evoked MFC connectivity pattern is positively correlated with behavior. Such results were not observed when applying the traditional univariate functional connectivity method.

### Study design considerations

First, previous studies have pointed out that neuroimaging findings often suffer from low reproducibility (Bennett & Miller, 2010; Noble et al., 2017; Poldrack et al., 2017). We tried to address this concern by seeking consistent findings across two datasets recorded at two different research sites using the same experimental paradigm. Furthermore, meta-analysis combining the two datasets helps to enhance statistical power when the need arises. Second, our paradigm included both instructional cues and choice cues (where subjects could choose where to attend). Given that instructional trials are twice as numerous as choice trials, we focused mainly on instructional trials in our main analysis. In Figure S1, however, we showed that similar results were observed for choice trials as well. This demonstrates that findings remained the same whether attention is deployed according to external instructions or purely internal decisions. Third, previous studies applying cross decoding techniques often included functional localizer trials, which are apart from the main attention or expectation paradigms and from which target classifiers are developed and applied to cross-decode cue-related activities. In our paradigm, the cue and the target occur during different task periods of the same trial, raising the concern that the cue evoked patterns may persist during target processing and thereby influence the cross-decoding results. We note, however, that both attended and ignored targets could follow an attention-directing cue, and that cue-based classifiers do not decode the neural patterns evoked by the ignored targets. This suggests that the cue evoked patterns do not contaminate the patterns evoked in the target period.

## Acknowledgements

This work was supported by National Institutes of Health grant MH117991 and National Science Foundation grant BCS-2318886 to G.R.M. and M.D.

