## Supplementary materials for "From Attention Control to Stimulus Selection: Neural Mechanisms Revealed by Multivariate Pattern and Functional Connectivity Analyses"

### *Example illustrating the Haufe et al. transformation*

To illustrate the idea, consider a two-voxel ROI where voxel 1 ($x_1(n)$) contains both the signal of interest, $s$(n), and noise, $d$(n), while voxel 2 ($x_2(n)$) contains only noise, $d$(n). A simple approach to recover the signal of interest is to compute the difference $x_1(n) - x_2(n) = \mathrm{w}^T\mathrm{x}(n)$, where $\mathrm{w} = [1, -1]^T$ is the weight vector of the model. However, this weight vector does not account for the physiological significance of the voxels, as voxel 2 does not contain information about the signal of interest and yet it is given the same weight as voxel 1. Assume that the signal of interest, $s$(n), is sampled from a normal distribution with a mean of 3 and a standard deviation of 1, while the noise, $d$(n), follows a normal distribution with a mean of 0 and a standard deviation of 1. The covariance matrix of the two-voxel system is: $\Sigma = \begin{bmatrix} 2.11 & 1.02 \\ 1.02 & 1.05 \end{bmatrix}$. By applying the Haufe et al. transformation, the resulting weight map is obtained as $[1.09, -0.03]^T$, which reflects the physiological significance of voxel 1 and discounts voxel 2.

### *Analysis of choice trials*

*Behavioral results:* See Table S1. (1) The UF dataset: for choose-left and choose-right trials, the target discrimination accuracy was 88.41 ± 1.88% and 90.78 ± 1.28% and the reaction time was 942.27 ± 47.75 ms and 938.25 ± 29.04 ms, respectively. There was no statistically significant difference between choose-left and choose-right trials in both target discrimination accuracy and reaction time ($p_{ACC} = 0.25$, $p_{RT} = 0.93$). (2) The UCD dataset: For choose-left and choose-right trials, the target discrimination accuracy was 84.07 ± 1.98% and 82.60 ± 2.14% and the reaction time was 1034.42 ± 48.06 ms and 1043.43 ± 45.84 ms, respectively. There was no statistically significant difference between choose-left and choose-right in both target discrimination accuracy and reaction time ($p_{ACC} = 0.34$, $p_{RT} = 0.72$).

| Dataset | Condition | Reaction Time (ms) | Accuracy (%) |
|---|---|---|---|
| UF | Cue Left | 918.56±24.49 | 85.72±2.14 |
| | Cue Right | 911.81±23.28 | 85.80±1.24 |
| | Choose Left | 942.27±47.75 | 88.41±1.88 |
| | Choose Right | 938.25±29.04 | 90.78±1.28 |
| UCD | Cue Left | 1027.02±55.60 | 82.45±2.30 |
| | Cue Right | 1041.87±54.95 | 81.97±2.10 |
| | Choose Left | 1034.42±48.06 | 84.07±1.98 |
| | Choose Right | 1043.43±45.84 | 82.60±2.14 |

Table S1. Behavioral results for choice trials.

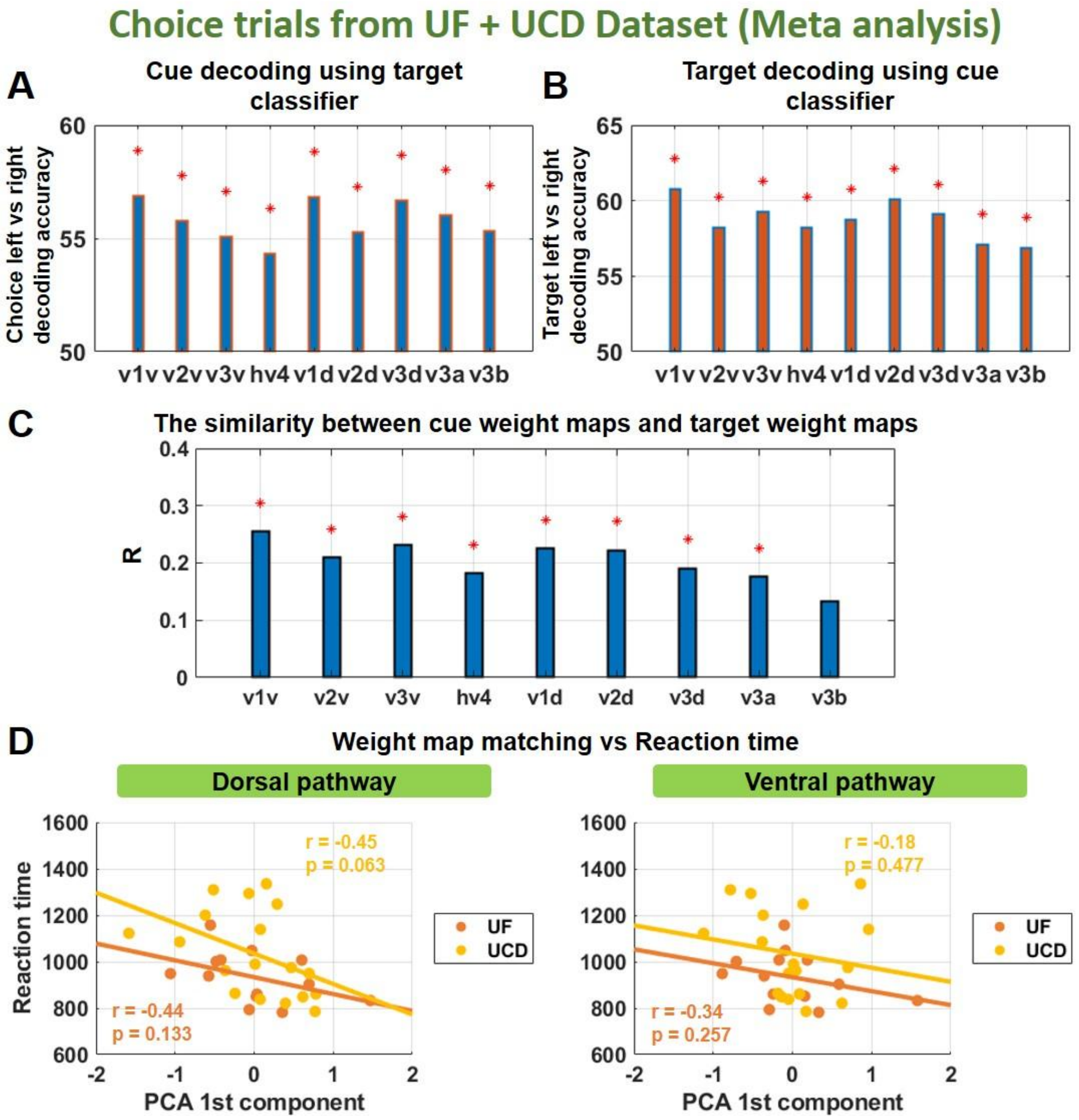


*Figure S1. Choice trial analysis. (A) Cross decoding analysis: decoding accuracies of choose left vs right using attended target-based classifiers. (B) Cross decoding analysis: decoding accuracies of attended left-target vs attended right-target using cue-based classifiers. (C) The correlation between cue weight maps and target weight maps for all participants and all ROIs. (D) Scatter plots showing relationship between the similarity of cue weight maps and target weight maps in the dorsal and ventral stream of visual cortex and the reaction time.*

*fMRI results:* In the main text we focused mainly on instructed trials because there are twice as many instructed trials as choice trials and it is known that larger amounts of data make machine learning based analysis more reliable. The choice trials, however, can offer two additional insights relative to instructed trials. First, a single choice cue results in two covert attentional states (attend left vs attend right), which helps to eliminate the concern that the pattern differences in early visual cortex are driven by the physical differences in the two attention-directing cues. Second, if the results we have reported so far hold for choice trials, then we can conclude that the same pattern matching mechanism governs the selection of the attended stimulus irrespective of whether the attentional states were generated via external instructions or internal decisions.

The same analysis as that applied to the instructed trials in the main text was applied to the choice trials. The results in Figure S1 are the meta-analysis results from combining the two datasets. In all ROIs, choice-left can be decoded from choice-right from cue-evoked data and attended left-target can be decoded from attended right-target from target-evoked data (not shown). Figure S1A and S1B show that for both target→cue cross decoding and cue→target cross decoding, the cross decoding accuracy is significantly above chance level in all ROIs, suggesting similarity between cue-related neural patterns and attended target-related patterns. This similarity was further quantified by the cue-target weight map correlation in Figure S1C. In all ROIs except v3b, the cue weight map and the target weight map exhibited significant positive correlations. Performing PCA analysis for weight map correlations in the dorsal and ventral pathways separately, and correlating the score on the first PCA component with the reaction time, we found that in the dorsal pathway, there was a significantly negative correlation with $r_{UF\ dorsal} = -0.44$ for the UF dataset, $r_{UCD\ dorsal} = -0.45$ for the UCD dataset, and the meta-analysis $p_{dorsal} = 0.03$; see Figure S1D. In the ventral pathway, however, no significant correlation was found, with $r_{UF\ ventral} = -0.34$, $r_{UCD\ ventral} = -0.18$, and the meta-analysis $p_{ventral} = 0.334$ (Figure S1D). Thus the results from the choice trials replicated the findings from the instructed trials.

***Multivariate functional connectivity (MFC) analysis of target processing***

We first tested whether attention modulated MFC. MFC was measured for the attended target and the ignored target separately and the results were averaged between the two hemispheres for each attention condition and compared across the two attention conditions. Among the 36 possible ROI pairs, Figure S2A listed the 19 connections whose MFC was enhanced by spatial attention ($p < 0.05$, FDR): v1v-hv4, v1v-v3a, v1v-v3d, v1v-v2d, v1v-v1d, v2v-hv4, v2v-v1d, v3v-v3d, v3v-v2d, v3v-v1d, hv4-v3a, hv4-v3d, hv4-v2d, v3a-v3d, v3a-v2d, v3a-v1d, v3d-v2d, v3d-v1d, and v2d-v1d. We then tested, for the attended target, whether MFC of these connections predicted behavior (Figure S2B). A negative correlation was found between the mean MFC connectivity and reaction time, where $r_{UF} = -0.30$, $r_{UCD} = -0.59$, and the meta-analysis $p = 0.018$, suggesting that the stronger the pattern-level interareal coordination in the visual cortex during attended target stimulus processing, the faster the reaction time. In contrast, comparing traditional univariate functional connectivity (UFC) between the attended and the ignored targets revealed no connections that were modulated by spatial attention (not shown).

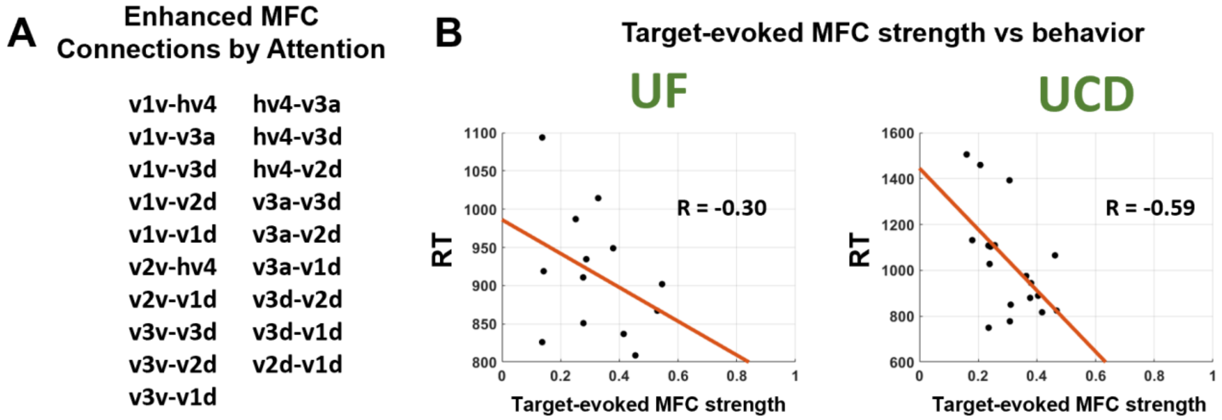


*Figure S2. Multivariate functional connectivity analysis of target-evoked activity. (A) Attention enhanced MFC for the following connections: v1v-hv4, v1v-v3a, v1v-v3d, v1v-v2d, v1v-v1d, v2v-hv4, v2v-v1d, v3v-v3d, v3v-v2d, v3v-v1d, hv4-v3a, hv4-v3d, hv4-v2d, v3a-v3d, v3a-v2d, v3a-v1d, v3d-v2d, v3d-v1d, and v2d-v1d. (B) The relationship between the mean MFC strength evoked by the attended targets and reaction time.*
